# Regulation of EGF-stimulated activation of the PI-3K/AKT pathway by exocyst-mediated exocytosis

**DOI:** 10.1101/2022.03.21.485168

**Authors:** Seong J. An, Alexander Anneken, Zhiqun Xi, Changseon Choi, Joseph Schlessinger, Derek Toomre

## Abstract

The phosphoinositide-3 kinase (PI-3K)/AKT cell survival pathway is an important pathway activated by EGFR signaling. Here we show, that in addition to previously described critical components of this pathway, i.e., the docking protein Gab1, the PI-3K/AKT pathway in epithelial cells is regulated by the exocyst complex, which is a vesicle tether that is essential for exocytosis. Using live-cell imaging, we demonstrate that PI(3,4,5)P_3_ levels fluctuate at the membrane on a minutes time scale and that these fluctuations are associated with local PI(3,4,5)P_3_ increases at sites where recycling vesicles undergo exocytic fusion. Supporting a role for exocytosis in PI(3,4,5)P_3_ generation, acute promotion of exocytosis by optogenetically driving exocyst-mediated vesicle tethering upregulates PI(3,4,5)P_3_ production and AKT activation. Conversely, acute inhibition of exocytosis using Endosidin2, a small-molecule inhibitor of the exocyst subunit Exo70, impairs PI(3,4,5)P_3_ production and AKT activation induced by EGF stimulation of epithelial cells. Moreover, prolonged inhibition of EGF signaling by EGFR tyrosine kinase inhibitors results in spontaneous reactivation of AKT without a concomitant relief of EGFR inhibition. However, this reactivation can be negated by acutely inhibiting the exocyst. These experiments demonstrate that exocyst-mediated exocytosis – by regulating PI(3,4,5)P_3_ levels at the plasma membrane – subserves activation of the PI-3K/AKT pathway by EGFR in epithelial cells.

## Introduction

Receptor tyrosine kinases (RTKs) are an important class of signal-transducing receptors that reside on the plasma membrane. By binding to extracellular ligands such as growth factors, RTKs initiate intracellular signaling cascades that regulate essential cellular processes, such as proliferation, cell survival, differentiation, migration, and metabolism (Lemmon and Schlessinger, 2010). Like most RTKs, the epidermal growth factor receptor (EGFR) undergoes ligand-induced dimerization, which activates the receptor through autophosphorylation of tyrosine residues within its cytoplasmic domain (Ullrich and Schlessinger, 1990). The resulting phosphotyrosines serve as docking sites to recruit cellular signaling molecules that activate multiple downstream signaling pathways including the phosphoinositide-3 kinase (PI-3K)/AKT and mitogen-activated protein kinase (MAPK) pathways (Sever and Brugge, 2015). Aberrant activation of these signaling pathways, through genetic alterations of RTKs or their downstream effectors, can lead to cancer and other diseases (Arteaga and Engelman, 2014).

The PI-3K/AKT pathway promotes cell survival, proliferation, growth, and metabolism through regulation of many downstream targets (Manning and Toker, 2017). PI-3 kinase is a heterodimer consisting of a regulatory subunit (p85) and a catalytic subunit (p110), whose enzymatic activity is promoted when the regulatory domain, through two SH2 domains, binds to phosphotyrosines in RTKs or associated docking proteins (Fruman et al., 2017). For EGF stimulation, PI-3 kinase activation primarily occurs through the docking protein Gab1 (Grb2-associated binder-1), which contains three canonical p85 binding motifs that are phosphorylated by EGFR (Mattoon et al., 2004). Activated PI-3 kinase then catalyzes phosphorylation of phosphatidylinositol 4,5-bisphosphate (PI(4,5)P_2_) to produce phosphatidylinositol 3,4,5-trisphosphate (PI(3,4,5)P_3_), which in turn can recruit AKT to the plasma membrane through its pleckstrin homology (PH) domain. At the membrane, AKT is then phosphorylated by other kinases to become activated (Manning and Toker, 2017).

It is well established that cell signaling by EGFR and other RTKs can be downregulated by receptor endocytosis (Lemmon and Schlessinger, 2010). Whether EGFR signaling is regulated by the converse membrane trafficking process, exocytosis, is undetermined. Exocytosis relies on vesicle tethers, which are protein complexes that bridge transport vesicles to their target membranes to facilitate intracellular membrane fusion (Hong and Lev, 2014). The exocyst is a type of multisubunit tether (Chou et al., 2016) that is essential for exocytosis of vesicles which recycle between the pericentriolar recycling endosome and the plasma membrane (Takahashi et al., 2005). It consists of eight subunits: Sec3, Sec5, Sec6, Sec8, Sec10, Sec15, Exo70 and Exo84. Three of these subunits have key known interactions with membranes. Exo70 and Sec3 (Zhang et al., 2008; He et al., 2007) both mediate vesicle tethering by interacting with PI(4,5)P_2_ on the plasma membrane, while Sec15 mediates attachment of the exocyst complex to the vesicle by binding the small GTPases Rab8 or Rab11 on the vesicle membrane (Wu et al., 2005). After a recycling vesicle is transported to the plasma membrane, the exocyst mediates an active tethering mechanism that promotes full vesicle fusion, leading to the complete merger of the vesicle with the membrane (An et al., 2021). Exocyst-mediated exocytosis is important for several cellular processes such as cell polarization (Polgar and Fogelgren, 2018), cell survival (Ahmed and Macara, 2017), migration (Rossé et al., 2006; Zou et al., 2006; Spiczka and Yeaman, 2008; Thapa et al., 2012) and glucose transport (Inoue, 2003).

In this paper, we demonstrate an unexpected relationship between RTK signaling and exocytosis. Based on our recent finding that optogenetically controlled exocyst-mediated exocytosis induces membrane expansion in cells (An et al., 2021), we investigated whether exocytosis upregulates PI(3,4,5)P_3_ as this phosphoinositide is known to play a key role in membrane remodeling (Burridge and Wennerberg, 2004). Here, we used a combination of approaches to test this hypothesis – including live-cell imaging, acute gain- and loss-function assays of the exocyst, and biochemical cell signaling assays. We show that exocyst-mediated exocytosis upregulates PI(3,4,5)P_3,_ and that this process underlies the activation of the PI-3K/AKT pathway downstream of EGF stimulation in epithelial cells.

## Results

### Exocytosis coincides spatiotemporally with an increase in PI(3,4,5)P_3_ level

Previously, we showed that vesicle tethering by the exocyst can be controlled by using a light-inducible heterodimerization system (An et al., 2021). This was achieved by first impairing membrane binding of the Exo70 exocyst subunit – by mutating two conserved lysines (K632A, K635A) near the C-terminal end that are critical for interactions with PI(4,5)P_2_ – and then attaching the photoreceptor cryptochrome 2 (CRY2) to the C-terminus of Exo70 so that membrane binding by the mutant Exo70 could be rescued by light-inducible heterodimerization of Exo70(K632A, K635A)-CRY2 with CIB1, the binding partner of CRY2, on the plasma membrane. One of the unexpected findings using this exocyst optogenetics system was that acute promotion of exocytosis causes the plasma membrane to expand, through formation and elongation of filopodia. Importantly, this membrane expansion was spatiotemporally associated with vesicle fusion events at the base of the expanding region. However, because the number of fusion events could not feasibly account for the extent of the membrane expansion, we reasoned that the membrane expansion could not merely reflect the delivery of membrane (i.e., lipids and proteins). Rather, we wondered whether exocytosis promoted membrane expansion by increasing levels of PI(3,4,5)P_3_ on the plasma membrane since this phosphoinositide is a key signaling molecule in membrane remodeling (Burridge and Wennerberg, 2004).

To test the hypothesis that exocytosis regulates PI(3,4,5)P_3_ levels, we explored PI(3,4,5)P_3_ dynamics at the membrane in live cells. When we transiently expressed the PI(3,4,5)P_3_ biosensor PH_AKT_-GFP (Várnai and Balla, 1998) at low levels in HeLa cells and imaged it using total internal reflection fluorescence (TIRF) microscopy, we found that the fluorescence of the biosensor at the membrane fluctuated on a minutes time scale (Fig. 1A, left). Such fluctuations were not seen when the biosensor was highly overexpressed (data not shown), or with CIB1-GFP-CAAX, a protein that is localized on the plasma membrane via a lipid anchor (Fig. 1A, right). The lack of fluctuations under these two latter conditions rules out the possibility that PH_AKT_-GFP fluctuations are due to undulations of the plasma membrane, i.e., changes in the proximity of the plasma membrane to the glass coverslip.

**Figure 1.**
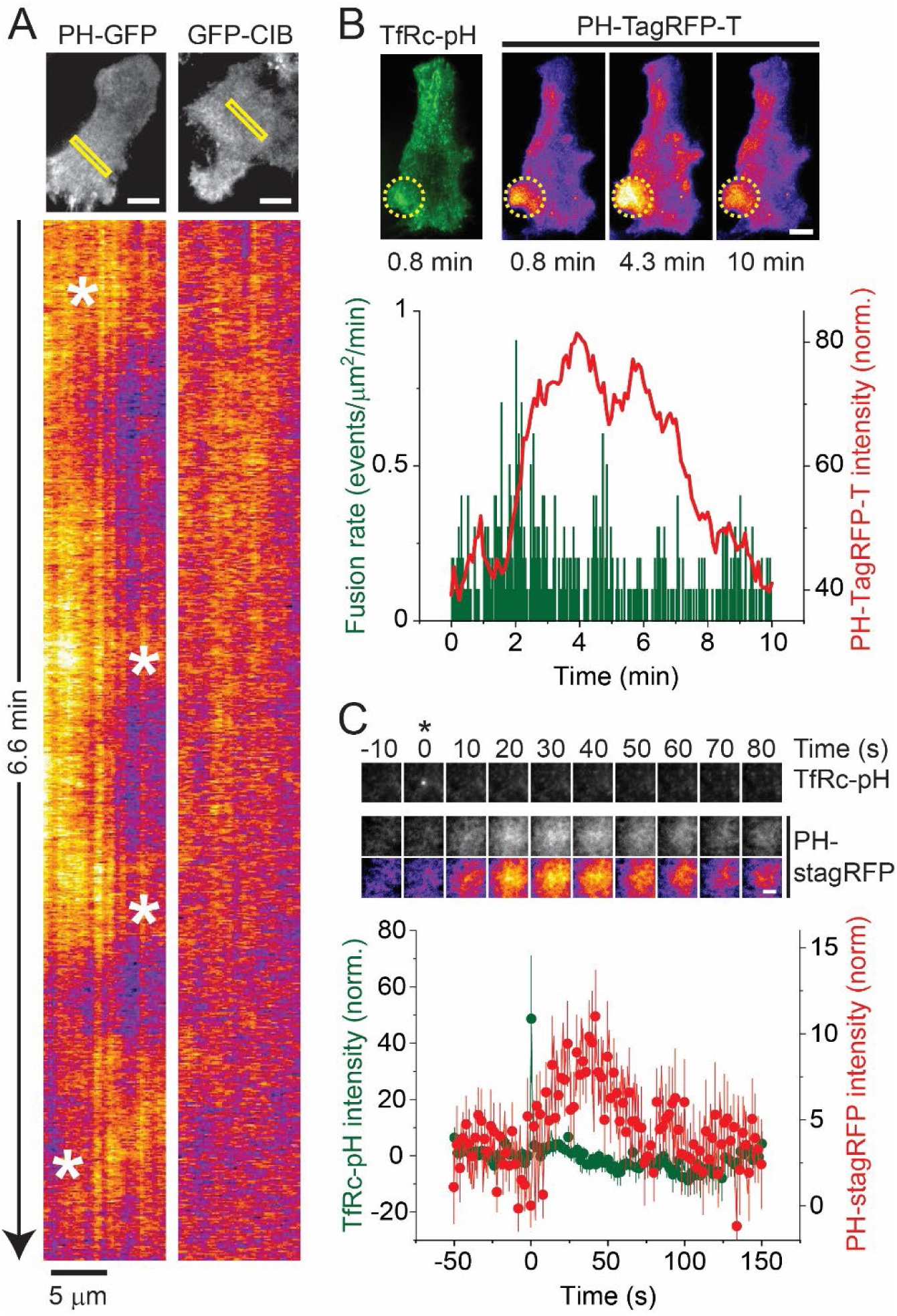
Exocytosis of recycling vesicles coincides spatiotemporally with an increase in PI(3,4,5)P_3_ level. (***A***) (*Upper*) TIRF micrographs showing the expression of PH_AKT_-GFP (*left*) or GFP-CIB-CAAX (*right*) in HeLa cells. (Scale bar: 20 µm.) (*Lower*) Kymographs of regions outlined by the yellow boxes in the upper panel showing fluorescence fluctuations (highlighted by white asterisks) with PH_AKT_-GFP but not with GFP-CIB-CAAX. Fluorescence is presented using a “fire” lookup table (ImageJ) to emphasize intensity changes from low (dark purple) to high (bright yellow). (***B***) (*Upper*) TIRF micrographs of TfRc-pH (green) and PH_AKT_-TagRFP-T (fire LUT) coexpressed in a HeLa cell. Dashed yellow circle highlights a fluctuation in the fluorescence of PH_AKT_-TagRFP-T that peaks at ∼4 min into the recording. (Scale bar: 20 µm.) (*Lower*) Plot showing time-dependent changes in the average intensity of PH_AKT_-TagRFP-T (red trace) and the fusion rate (10-s bins; green bars) in the region outlined by the dashed yellow circle in the upper panels. (***C***) (*Upper*) Average TIRF image sequence of vesicles undergoing exocytosis (n = 31 events), time aligned to the moment of fusion (t = 0, asterisk). Images of PH_AKT_-stagRFP are also displayed using a fire LUT to emphasize confinement of the fluorescence increase to the site of vesicle fusion. (Scale bar: 2 µm.) (*Lower*) Average intensity traces of PH_AKT_-stagRFP (red) and TfRc-pH (green), time aligned to the moment of fusion. Data are shown as mean± S.E.M.

To test whether PH_AKT_-GFP fluctuations coincide with exocytosis, we performed two-color imaging experiments with the pH-sensitive exocytosis reporter transferrin receptor-pHluorin (TfRc-pH; Merrifield et al, 2005), which emits in the green channel (∼509 nm), and the PI(3,4,5)P_3_ biosensor with its GFP replaced by TagRFP-T, a fluorescent protein that emits in the red channel (∼584 nm; Shaner et al., 2008). Figure 1B shows an example of such a two-color movie. Here, an initial rise in PH_AKT_-TagRFP-T fluctuation coincided with a burst of exocytic fusion events. Notably, the duration of the PH_AKT_-TagRFP-T fluctuation lasted for several minutes, while the elevation in the fusion rate quickly subsided after momentarily increasing by ∼4-fold. This difference in duration between PH_AKT_-TagRFP-T fluctuation and fusion rate increase was recapitulated at the level of a single vesicle: When a brighter version of TagRFP-T (Mo et al., 2020) was attached to the PI(3,4,5)P_3_ biosensor (PH_AKT_-stagRFP), its signal increased as a cloud of fluorescence that persisted for over ∼1 min after an exocytic event (Fig. 1C). Evidently, an increase in PH_AKT_-stagRFP fluorescence can be spatially confined at the site of vesicle fusion, long after a vesicle has fused. These results suggest that vesicle fusion locally generates PI(3,4,5)P_3_ and that multiple vesicles can sustain a longer lasting PI(3,4,5)P_3_ fluctuation by undergoing exocytosis around the same time and place.

### Acute promotion of exocytosis upregulates PI(3,4,5)P_3_

We wished to test whether acutely promoting exocytosis can cause an increase in PI(3,4,5)P_3_ levels. To this end, we transiently coexpressed Exo70(K632A, K635A)-CRY2-mCherry and plasma-membrane targeted CIB1-CAAX to control exocytosis optogenetically (Fig. 2A) and PH_AKT_-GFP to visualize PI(3,4,5)P_3_. Illumination of cells using 100-ms pulses of 488-nm light (to activate CRY2 and image PI(3,4,5)P_3_ simultaneously) caused a steady rise in PH_AKT_-GFP fluorescence that plateaued after ∼5 min (Fig. 2B-D). Consistent with the fluorescence increase reflecting upregulation of PI(3,4,5)P_3_, a much smaller increase was seen when we imaged a version of PH_AKT_-GFP carrying a mutation (R25C) that has greatly diminished affinity for PI(3,4,5)P_3_ (Franke et al., 1997). To assess the relative magnitude of PI(3,4,5)P_3_ generation by exocyst optogenetics, we next imaged PI(3,4,5)P_3_ during EGF stimulation (10 ng/mL) under the same imaging conditions. Interestingly, while EGF stimulation caused PH_AKT_-GFP fluorescence to rise ∼2-fold higher compared to exocyst optogenetics, the fluorescence peaked earlier (∼2 min) and returned to baseline, unlike with exocyst optogenetics where it remained elevated. \Again, a much smaller increase in PH_AKT_-GFP fluorescence was seen with the R25C mutant of the PI(3,4,5)P_3_ biosensor in EGF-stimulated cells compared to the wildtype sensor.

**Figure 2.**
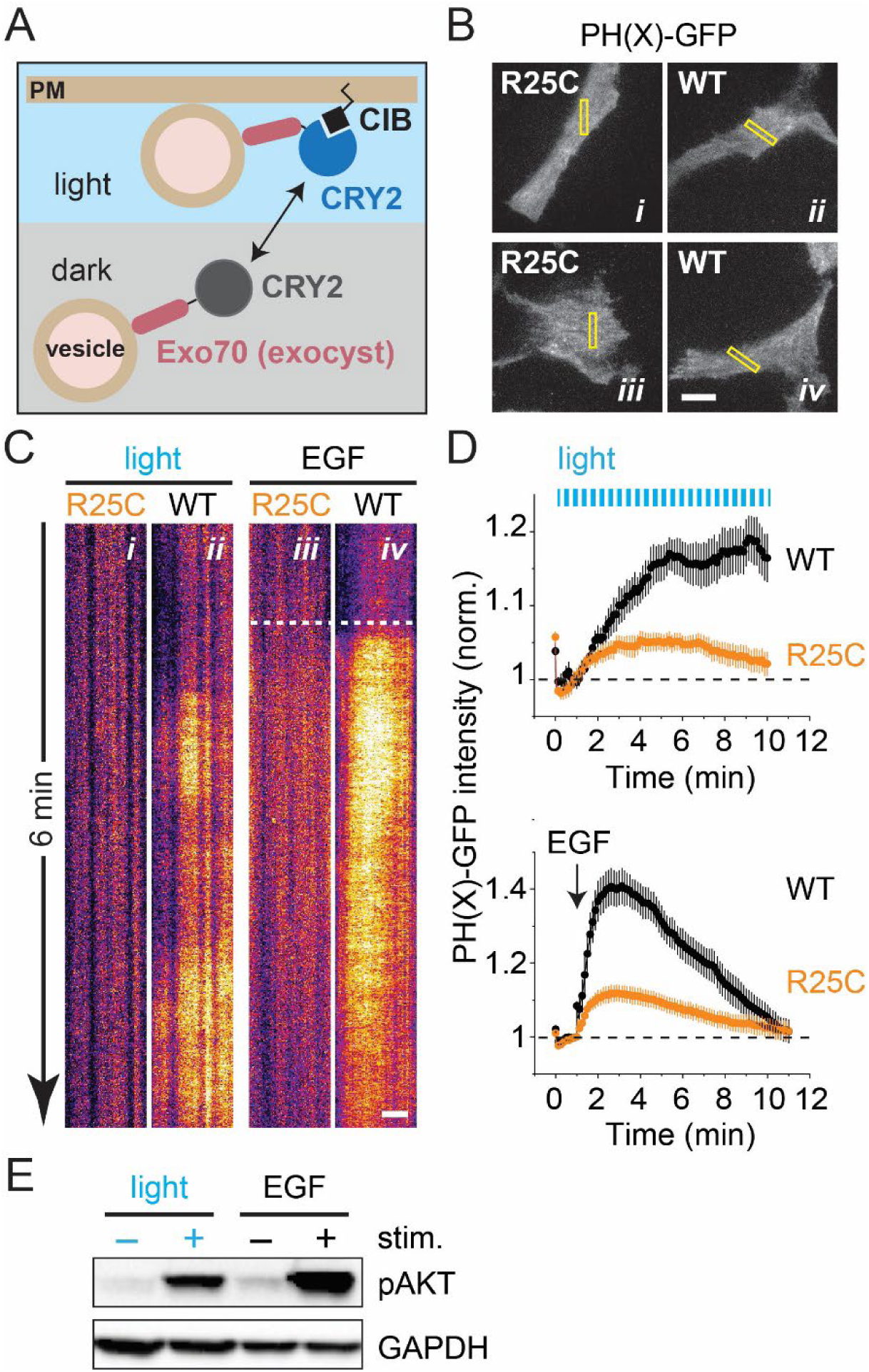
Acute promotion of exocyst-mediated exocytosis upregulates PI(3,4,5)P_3_ and activates AKT. (***A***) Schematic drawing depicting optogenetic promotion of vesicle tethering by light-induced heterodimerization of Exo70(K632A, K635A)-CRY2-mCherry on the vesicle and CIB-CAAX on the plasma membrane (PM). Note that only Exo70 within the exocyst complex is depicted for clarity. (***B***) TIRF micrographs of wildtype or the R25C mutant of PH_AKT_-GFP expressed in HeLa cells before stimulation with 488-nm light (panels *i* and *ii*) or EGF (panels *iii* and *iv*). Note that the cells for optogenetic stimulation also coexpress Exo70(K632A, K635A)-CRY2-mCherry and CIB-CAAX. (Scale bar: 20 µm.) (***C***) Kymographs of regions outlined by the yellow boxes in *B* showing fluorescence changes induced by 100-ms pulses (2 Hz) of 488-nm light or EGF (10 ng/mL). Fluorescence is presented using a “fire” lookup table (ImageJ). Dashed white line in the right two kymographs indicates start of EGF stimulation. (Scale bar: 10 µm.) (***D***) Time-dependent changes in the average fluorescence (of the whole membrane) for wildtype PH_AKT_-GFP (n = 11 cells) and the R25C mutant of PH_AKT_-GFP (n = 10 cells) during stimulation with 488-nm light (upper plot) or for wildtype PH_AKT_-GFP (n = 12 cells) and the R25C mutant of PH_AKT_-GFP (n = 13 cells) during stimulation with EGF (lower plot). Data are shown as mean± S.E.M. Note that traces are normalized to prestimulation intensity (dashed lines). (***E***) Comparison of AKT activation by exocyst optogenetics and EGF stimulation. HeLa cells were grown on 35-mm dishes and stimulated with blue light on an LED array or with EGF (100 ng/mL) for 5 min at 37 °C. Cell lysates were subjected to sodium dodecyl sulfate polyacrylamide gel electrophoresis (SDS/PAGE) and analyzed for activation of AKT by immunoblotting with antibodies for pAKT (Ser473).

If exocyst optogenetics generates PI(3,4,5)P_3,_ as the above imaging experiments suggest, then it might affect cell signaling activated by PI(3,4,5)P_3_, such as the AKT pathway. To test this, we performed whole-dish optogenetics experiments to check for AKT activation using biochemical techniques. HeLa cells were transiently transfected with Exo70(K632A, K635A)-CRY2-mCherry and CIB1-CAAX and then optically stimulated *en masse* with a light-emitting diode array for 5 min at 37 °C. After stimulation, cell lysates were subjected to SDS/PAGE analysis followed by immunoblotting with anti-phospho-AKT antibodies. Figure 2E shows that exocyst optogenetics could indeed induce AKT phosphorylation, although to a lesser extent than EGF stimulation (100 ng/mL, 5 min at 37 °C). While this difference is consistent with the greater upregulation of PI(3,4,5)P_3_ by EGF in the imaging experiments described in Figures 2B to D, it should be noted that AKT phosphorylation in the whole-dish optogenetics experiment was likely suboptimal given that only a fraction of cells are adequately transfected with all components of the optogenetics system. Notwithstanding this point, the above results demonstrate that acute promotion of exocytosis not only upregulates PI(3,4,5)P_3_ but also activates AKT.

### Acute inhibition of exocytosis downregulates PI(3,4,5)P_3_

We reasoned that if acute promotion of exocytosis upregulates PI(3,4,5)P_3_, then the converse experiment, acute inhibition of exocytosis, might have the opposite effect. To explore this possibility, we performed a series of experiments with Endosidin2 (Es2), a small molecule inhibitor of the exocyst (Fig. 3A; Zhang et al., 2016). Mechanistically, Es2 binds to Exo70, which inhibits not only Exo70 binding to the plasma membrane but also transferrin recycling in HeLa cells. We confirmed the inhibitory effect of Es2 on exocytosis by performing TIRF microscopy of exocytosis, again by imaging TfRc-pH in HeLa cells, and measuring the fusion rate before and after a 30-min treatment with increasing concentrations of the drug. We found that Es2 inhibited exocytosis in a dose-dependent manner, with an IC_50_ of 25.8 ± 7.3 µM (n = 5; Fig. 3B), consistent with the reported micromolar affinity of Es2 for Exo70 (Zhang et al., 2016). It should be noted that maximal inhibition of exocytosis was only 49.2 ±7.3% at the highest concentration tested. However, this is consistent with the ∼50% colocalization of TfRc-pH-containing vesicles with the exocyst (Rivera-Molina and Toomre, 2013).

**Figure 3.**
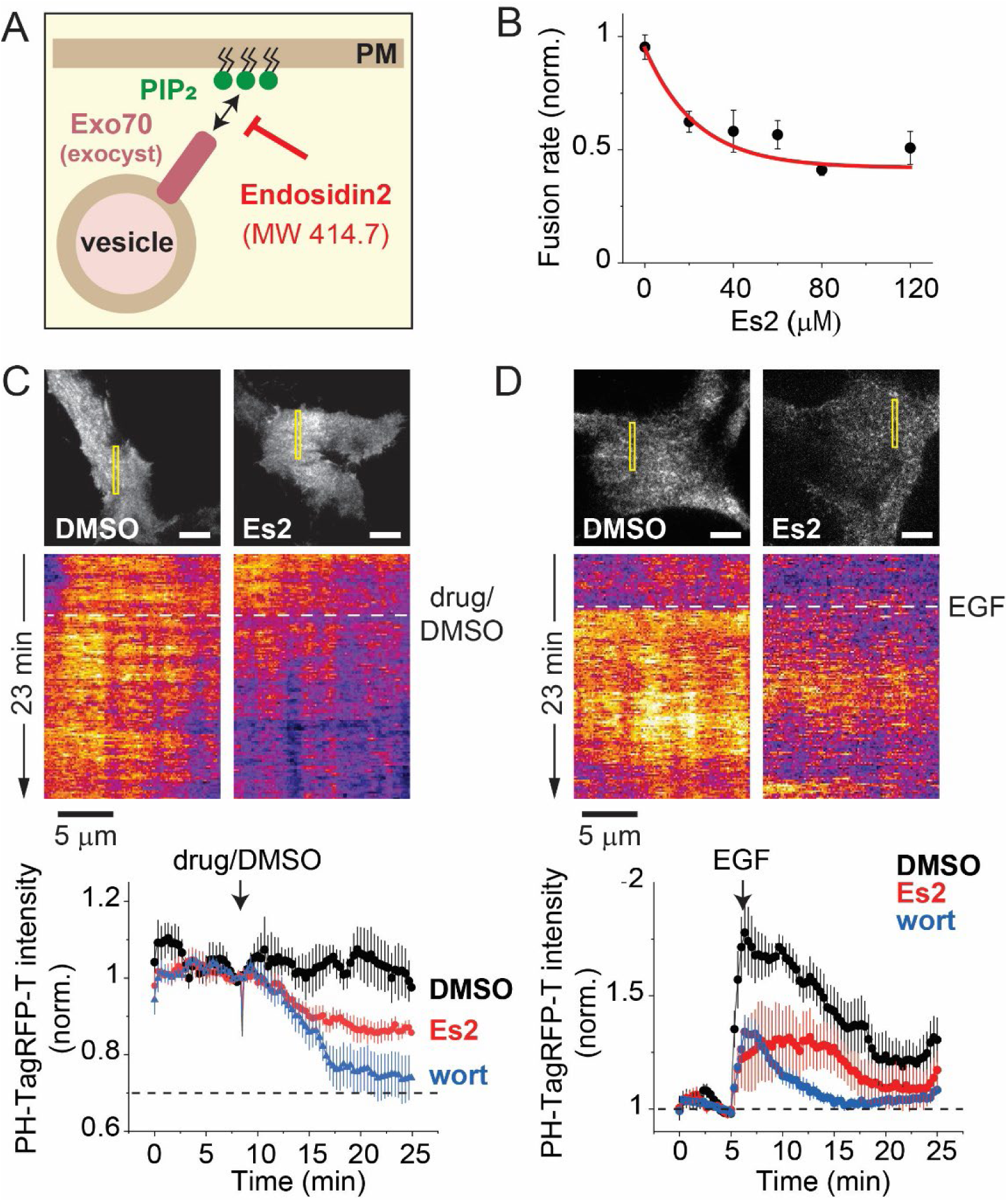
Acute inhibition of exocyst-mediated exocytosis downregulates PI(3,4,5)P_3_. (***A***) Schematic drawing depicting inhibition of vesicle tethering by Endosidin2 (Es2), via inhibition of Exo70 binding to PI(4,5)P_2_ on the plasma membrane. Note that only Exo70 within the exocyst complex is depicted for clarity. (***B***) Inhibition of exocyst-mediated exocytosis by Es2. HeLa cells expressing TfRc-pH were imaged for 5 min, then treated with 0 µM (DMSO; n = 5 cells), 20 µM (n = 5 cells), 40 µM (n = 3 cells), 60 µM (n = 5 cells, 80 µM (n = 4 cells) or 120 µM (n = 4 cells) Es2 for 30 min at 37 °C, and finally re-imaged for 5 min to record exocytosis, before and after drug treatment. The fusion rate (events/µm^2^/min) was calculated for each period and the normalized rate change (post/pre) for each experiment was plotted against Es2 concentration. Datapoints were fitted (red) using a dose-response curve fitting function (Origin). (***C***) The effect of Es2 on basal PI(3,4,5)P_3_ levels. (*Upper*) TIRF micrographs of PH_AKT_-TagRFP-T in HeLa cells before treatment with either DMSO or Es2 (80 µM). (Scale bar: 20 µm.) (*Middle*) Kymographs of regions outlined by the yellow boxes in the upper panels. White dashed line indicates start of DMSO or Es2 treatment. Fluorescence is presented using a “fire” lookup table (ImageJ). (*Lower*) Time-dependent changes in PH_AKT_-TagRFP-T fluorescence caused by the addition of DMSO (n = 4 cells), Es2 (n = 5 cells) and wortmannin (100 nM; n = 6 cells). Data are shown as mean± S.E.M. Note that the fluorescence decrease does not exceed 30% (dashed line), even for wortmannin. (***D***) The effect of Es2 pretreatment (30 min at 37 °C) on EGF-stimulation of PI(3,4,5)P_3_ production. (*Upper*) TIRF micrographs of PH_AKT_-TagRFP-T in HeLa cells before treatment with either DMSO or Es2 (80 µM). (Scale bar: 20 µm.) (*Middle*) Kymographs of regions outlined by the yellow boxes in the upper panels. White dashed line indicates start of EGF (10 ng/mL) stimulation. Fluorescence is presented using a “fire” lookup table (ImageJ). (*Lower*) Time-dependent changes in PH_AKT_-TagRFP-T fluorescence during EGF stimulation in cells pretreated with DMSO (n = 11 cells), Es2 (n = 13 cells) and wortmannin (100 nM; n = 14 cells). Data are shown as mean± S.E.M. Note that traces are normalized to prestimulation intensity (dashed line).

Next, we used Es2 to test the effect of inhibiting exocytosis on basal PI(3,4,5)P_3_ levels. There were several considerations in designing these experiments. First, since not all cells are typically “active” vis-à-vis exocytosis (our observations), we selected cells that showed robust TfRc-pH fusion activity. Second, because it was necessary to image PI(3,4,5)P_3_ for tens of minutes to allow drug diffusion into cells, we used a slower acquisition rate (0.2 Hz compared to 2Hz for imaging exocytosis) to avoid photobleaching of PH_AKT_-TagRFP-T. Finally, after recording a sufficiently long (∼8.3 min) baseline signal, we added either Es2 (80 µM), the PI-3 kinase \inhibitor wortmannin (100 nM) as a positive control or the drug vehicle DMSO as a negative control. The results presented in Figure 3C show that Es2 caused a time-dependent decrease in PH_AKT_-TagRFP-T fluorescence that was not seen with DMSO (13.0 ± 1.6% vs. −0.4 ± 5.8%, P = 0.042) but somewhat smaller than that seen with wortmannin (13.0 ± 1.6% vs. 26.2 ± 5.9%, P = 0.082). It should be noted that the fluorescence decrease was less than 30% (dashed line in lower plot of Fig. 3C) even with wortmannin because the penetration depth of the evanescent field is greater than the thickness of the membrane (∼100 nm vs. ∼10 nm), which allows for much of the fluorescence to be derived from PH_AKT_-TagRFP-T molecules in the cytosol, not just at the membrane.

The above experiments demonstrate that exocytosis contributes to steady-state levels of PI(3,4,5)P_3_. To test whether exocytosis is involved in upregulation of PI(3,4,5)P_3_ by RTK signaling, we performed EGF stimulation experiments with cells that were starved overnight in serum-free media. As before, we selected cells based on the presence of fusion events. We then treated the cells for 30 min with either Es2 (80 µM), wortmannin (100 nM) or DMSO for 30 min at 37 °C before stimulating them with EGF (10 ng/mL). Figure 3D shows that Es2 pretreatment caused an overall increase in PH_AKT_-TagRFP-T fluorescence that was smaller compared to DMSO (integrated signal of 24.6 ± 10.2 a.u. vs. 53.8 ± 7.2 a.u., P = 0.034), but somewhat larger than the overall increase after wortmannin pretreatment (24.6 ± 10.2 a.u. vs. 13.2 ± 2.3 a.u., P = 0.27). These results demonstrate that exocytosis contributes to PI(3,4,5)P_3_ upregulation by EGF stimulation.

### Acute inhibition of exocytosis inhibits AKT stimulation in epithelial cells

The data presented so far suggest that (*i*) exocytosis of individual vesicles generates PI(3,4,5)P_3_ (Fig. 1C); (*ii*) acute optogenetic promotion of exocyst-mediated exocytosis upregulates PI(3,4,5)P_3_ (Fig. 2B-D) and AKT phosphorylation (Fig. 2E); and (*iii*) acute inhibition of exocyst-mediated exocytosis using Es2 downregulates steady-state (Fig. 3C) and EGF-stimulated PI(3,4,5)P_3_ levels (Fig. 3D). As such, we tested whether Es2 inhibits AKT activation by EGF stimulation. HeLa cells were starved in serum-free media and then treated with increasing concentrations of Es2 for 30 min at 37 °C before stimulation with EGF (100 ng/mL) for 5 min. Cell lysates were then subjected to immunoblotting with anti-pAKT antibodies, as well as anti-pMAPK antibodies, to monitor AKT and MAPK stimulation, respectively. Surprisingly, we found that Es2 caused very weak inhibition of AKT phosphorylation (Fig. 4A). One possible explanation for this result is that PI(3,4,5)P_3_ upregulation by *endogenous* exocytosis is not tightly coupled to AKT stimulation in HeLa cells. Because it has been reported that shRNA-mediated knockdown of the exocyst subunit Sec8 or Sec10 significantly reduces phospho-AKT levels in murine mammary gland NMuMG cells (Ahmed and Macara, 2017), we wondered whether the upregulation of PI(3,4,5)P_3_ by exocyst-mediated exocytosis is coupled to AKT stimulation specifically in epithelial cells.

**Figure 4.**
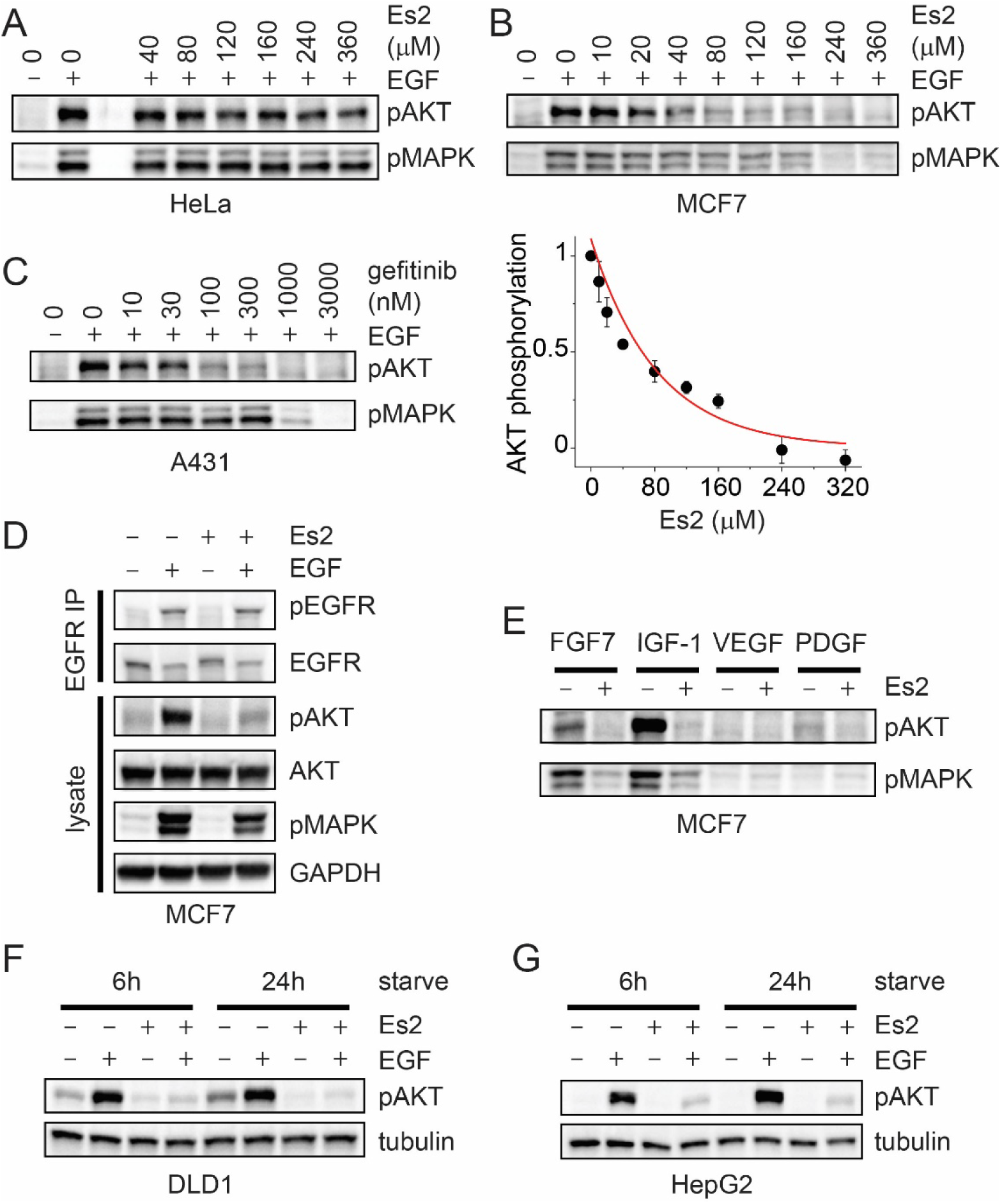
The effects of Es2 on EGF signaling in different cell lines. (***A***) HeLa cells were cultured for 1 d, starved for 1 d, treated with increasing concentrations of Es2 (0−360 µM) for 30 min at 37 °C, and then stimulated with EGF (100 ng/mL) for 5 min. Cell lysates were subjected to SDS/PAGE and analyzed for activation of AKT and MAPK by immunoblotting with antibodies for pAKT (Ser473) and pMAPK (Thr202, Tyr204), respectively. (***B***) (*Upper*) MCF7 cells were cultured for 7 d, starved for 2 d, treated with increasing concentrations of Es2 (0−360 µM) for 30 min at 37 °C, and then stimulated with EGF (100 ng/mL) for 5 min. AKT and MAPK activation were analyzed as above. (*Lower*) Analysis of the dose-dependent inhibition of AKT phosphorylation by Es2 (n = 3 cells). Data are shown as mean± S.E.M. Datapoints were fitted (red) using a dose-response curve fitting function (Origin). (***C***) A431 cells were cultured for 3 d, starved for 6 h, treated with increasing concentrations of gefitinib (0−3 µM) for 30 min at 37 °C, and then stimulated with EGF (10 ng/mL) for 5 min. AKT and MAPK activation were analyzed as above. (***D***) MCF7 cells were cultured for 7 d, starved for 2 d, treated with Es2 (240 µM) for 30 min at 37 °C, and then stimulated with EGF (100 ng/mL) for 5 min. EGFR was immunoprecipitated from cell extracts by using anti-EGFR antibodies and EGFR stimulation and expression were analyzed by immunoblotting the precipitates with anti-pEGFR and anti-EGFR antibodies, respectively. AKT and MAPK activation were analyzed as above. (***E*** and ***F***). DLD1 cells and HepG2 cells were cultured for 4 d or 5 d, respectively, starved for two different durations (6 and 24 h), treated with Es2 (360 µM) for 30 min at 37 °C, and then stimulated with EGF (100 ng/mL) for 5 min. AKT and MAPK activation were analyzed as above.

To test this notion, we turned to MCF7 cells, which are a human mammary epithelial cell line. MCF7 cells were cultured for 7 d (first 3 d with 0.1 mg/mL insulin) to near confluence (∼80%), starved in serum-free media (0.2% BSA) for 2 d and then treated with increasing concentrations of Es2 for 30 min at 37 °C before stimulation with EGF (100 ng/mL) for 5 min. Cell lysates were then subjected to immunoblotting with anti-pAKT antibodies and anti-pMAPK antibodies to monitor AKT and MAPK stimulation, respectively. The results presented in Figure 4B show that Es2 completely inhibited AKT phosphorylation in a dose-dependent manner with an IC_50_ of 60.7 ± 6.1 µM (n = 4), demonstrating that EGF-induced AKT stimulation can indeed be dependent on exocyst-mediated exocytosis.

In MCF7 cells, Es2 also inhibited phosphorylation of MAPK, although higher concentrations of the drug were needed to inhibit MAPK stimulation than AKT stimulation (Fig. 4B). A dependence of MAPK stimulation on PI-3 kinase has been reported in MCF7 cells (Ebi et al., 2013). Interestingly, we found that the differential sensitivity of AKT and MAPK stimulation to Es2 was similar to that of the EGFR tyrosine kinase inhibitor gefitinib (Fig. 4C), which also inhibited MAPK stimulation at higher concentrations than AKT stimulation in another epithelial cell line, A431 cells. These results suggest that if exocyst-mediated exocytosis regulates EGFR signaling, then it might do so by regulating a signaling node close to the activation of EGFR itself. Thus, a simple but possibly trivial explanation for the ability of Es2 to inhibit AKT and MAPK stimulation is that Es2 inhibits EGFR activation. To test this possibility, we examined the phosphorylation of EGFR after Es2 treatment. We repeated the previous experiment but this time, after EGF stimulation, we immunoprecipitated EGFR from cell extracts by using anti-EGFR antibodies and monitored EGFR stimulation and expression by immunoblotting the precipitates with anti-pEGFR and anti-EGFR antibodies, respectively. Immunoprecipitation of EGFR was necessary since MCF7 cells express very low levels of the receptor, which precludes its detection by immunoblotting of cell lysates. As shown in Figure 4D, Es2 (240 µM) strongly inhibited AKT phosphorylation without inhibiting EGFR phosphorylation or affecting EGFR levels. These results demonstrate that Es2 inhibits AKT and MAPK stimulation downstream of EGFR activation.

Next, we tested whether inhibition of exocyst-mediated exocytosis by Es2 affects AKT stimulation in two other epithelial cell lines: human colorectal DLD1 cells and human hepatoma HepG2 cells. DLD1 and HepG2 cells were cultured for 4 d to confluence or 3 d to 80% confluence, respectively, then starved in serum-free media (for 6 or 24 h) and treated with Es2 (320 µM) for 30 min before stimulation with EGF (100 ng/mL) for 5 min. Cell lysates were then subjected to immunoblotting with anti-pAKT antibodies to monitor AKT stimulation. Figures 4E and 4F show that Es2 strongly inhibited AKT phosphorylation in both cell lines, suggesting that exocyst-mediated exocytosis is coupled to EGF-induced AKT stimulation in different types of epithelial cells.

### Es2 inhibits a novel form of drug resistance in A431 cells

In the next set of experiments, we studied a fourth epithelial cell line, human epidermoid A431 cells. These cells have high levels of EGFR expression, making it convenient to monitor the stimulation and expression of EGFR by immunoblotting cell lysates. We first tested whether A431 cells are sensitive to Es2 regarding EGF-induced AKT stimulation. A431 cells were cultured for 3 d until ∼80% confluent, starved in serum-free media (for 6 or 24 h) and treated with Es2 (320 µM) for 30 min before stimulation with EGF (10 ng/mL) for 5 min. Cell lysates were then subjected to immunoblotting with anti-pAKT antibodies and anti-pMAPK antibodies to monitor AKT and MAPK stimulation, respectively. As with the other epithelial cell lines, Es2 strongly inhibited both AKT and MAPK stimulation (Fig. 5A), without affecting EGFR activation (Supplementary Fig.1). Interestingly, however, in A431 cells (Fig. 5A) – but less so in DLD1 (Fig. 4F) or HepG2 cells (Fig. 4G) – both basal and EGF-stimulated AKT but not MAPK phosphorylation was greater after 24 h starvation than with 6 h starvation. This suggests that suppressing cell signaling in A431 cells sensitized AKT but not MAPK stimulation.

**Figure 5.**
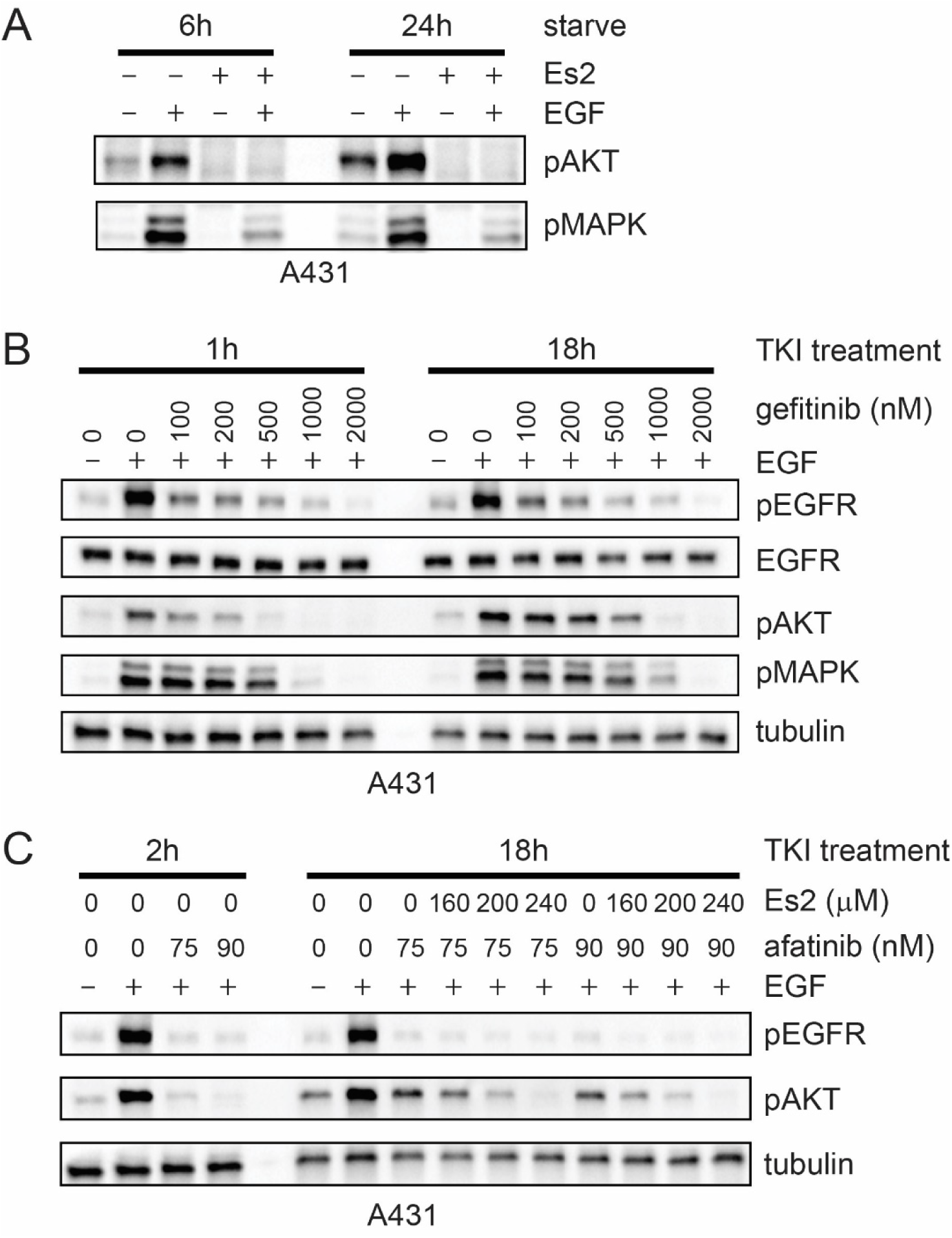
Es2 inhibits a novel form of drug resistance in A431 cells. (***A***) A431 cells were cultured for 3 d, starved for either 6 or 24 h, treated with Es2 (320 µM) for 30 min at 37 °C, and then stimulated with EGF (10 ng/mL) for 5 min. Cell lysates were subjected to SDS/PAGE and analyzed for activation of AKT and MAPK by immunoblotting with antibodies for pAKT (Ser473) and pMAPK (Thr202, Tyr204), respectively. (***B***) A431 cells were cultured for 3 d, starved for 6 h, then treated with increasing concentrations of gefitinib (0−2 µM) for two different durations (1h and 18 h) at 37 °C, and finally stimulated with EGF (10 ng/mL) for 5 min. (***C***) A431 cells were cultured for 3 d, starved for 6 h, then treated with the indicated concentrations of afatinib for two different durations (2h and 18 h) at 37 °C. Immediately after each afatinib treatment period, the cells were treated with the indicated concentrations of Es2 for 30 min at 37 °C, and then stimulated with EGF (10 ng/mL) for 5 min.

One explanation for the above finding is that suppressing cell signaling causes a compensatory adaptation that upregulates AKT activation. To explore this idea, we examined the effect of EGFR inhibitors on AKT stimulation over time. A431 cells were cultured for 3 d, starved for 6 h and then treated with increasing concentrations of the EGFR TKI gefitinib to inhibit EGF signaling for either 1 or 18 h. Immediately after each TKI treatment period, the cells were stimulated with EGF (10 ng/mL) for 5 min. The results presented in Figure 5B show that AKT phosphorylation was strongly inhibited in a dose-dependent manner after 1 h of TKI treatment. As in an earlier experiment using a short treatment period (30 min; Fig. 4C), AKT and MAPK inhibition displayed different dose responses to gefitinib, with MAPK inhibition requiring higher drug concentrations. However, after the 18 h treatment period, AKT phosphorylation levels rose again at submaximal gefitinib concentrations (< 2 µM), shifting the dose response of AKT inhibition to the right so that it looked similar to the dose response of MAPK inhibition. Importantly, this change occurred without a commensurate change in the dose response of EGFR inhibition to gefitinib. Thus, A431 cells rapidly developed resistance to gefitinib with respect to AKT stimulation.

Finally, we wondered whether the re-emergence of AKT stimulation during prolonged EGFR inhibition depended on exocyst-mediated exocytosis. This could be tested by treating cells first with an EGFR TKI to inhibit EGFR and then with Es2 to inhibit exocytosis. Initial experiments indicated possible drug-drug interactions between gefitinib and Es2, which necessitated sequential but separate drug treatments. As such, we turned to a different EGFR TKI, afatinib. Unlike gefitinib, this TKI irreversibly inhibits EGFR, allowing us to add Es2 separately after TKI treatment without potentially reversing EGFR inhibition. As in the previous experiment, A431 cells were cultured for 3 d and starved for 6 h. The cells were then treated with afatinib at two different concentrations (75 and 90 nM) to ensure sufficient but submaximal AKT inhibition for either 2 or 18 h. Immediately after each TKI treatment period, the cells were treated with Es2 at three ascending concentrations (160, 200 and 240 µM) and then finally stimulated with EGF (10 ng/mL) for 5 min. The results presented in Figure 5C show that AKT phosphorylation by EGF was inhibited down to pre-stimulation levels during the first 2 h of afatinib treatment but reappeared after 18 h even though EGFR phosphorylation remained low. Importantly, subsequently treating cells with Es2 after 18 h strongly inhibited the reappearance of phosphorylated AKT in a dose-dependent manner. Thus, these results demonstrate that reactivation of AKT during prolonged EGFR inhibition with TKIs can be negated by inhibiting exocyst-mediated exocytosis.

## Discussion

The PI-3/AKT pathway is one of the main cellular signaling pathways that are activated by RTKs and other cellular cues to mediate a critical anti-apoptotic, i.e., cell survival signal. While several RTKs, e.g., PDGFR and ErbB3, activate PI3-kinase through formation of direct contacts between the regulatory p85 subunit of PI-3 kinase with specific tyrosine phosphorylation sites on activated RTK, other RTKs, e.g., EGFR and INSR, utilize closely associated tyrosine phosphorylated docking proteins such as Gab1 and IRS1, respectively, for p85 recruitment and PI-3 kinase activation. EGFR utilizes an additional indirect mechanism for PI-3 kinase recruitment and activation through EGF-induced heterodimerization with ErbB3 resulting in tyrosine phosphorylation of several specific docking sites for PI-3 kinase in the cytoplasmic domain of ErbB3. It is noteworthy that both Gab1 and AKT utilize their PH domains for translocation to the cell membrane by specific binding to PI(3,4,5)P_3_ molecules; membrane translocation of Gab1 and AKT is an essential step in a highly regulated EGF-induced positive feedback mechanism of PI-3K/AKT activation (Rodrigues et al., 2000). In this report, we present several lines of evidence that exocyst-mediated exocytosis regulates AKT stimulation downstream of EGFR signaling primarily in epithelial cells. First, our live-cell imaging demonstrates that when an individual recycling vesicle undergoes exocytosis, the local level of PI(3,4,5)P_3_ increases for ∼1 min. And when multiple vesicles undergo exocytosis around the same time, a longer lasting PI(3,4,5)P_3_ fluctuation occurs at the membrane. We then demonstrate that exocytosis and PI(3,4,5)P_3_ levels are causally related by controlling the exocyst through acute gain- and loss-of-function methods. We show that acutely promoting exocytosis through exocyst optogenetics upregulates PI(3,4,5)P_3_ and AKT phosphorylation to an extent comparable to that of EGF stimulation. Conversely, we show that acutely inhibiting exocyst-mediated exocytosis using Es2, a small-molecule inhibitor of Exo70, downregulates PI(3,4,5)P_3_ levels to an extent that is comparable to that by the PI-3 kinase inhibitor wortmannin. Collectively, these imaging data not only support our biochemical experiments showing that exocyst-mediated exocytosis regulates AKT stimulation, but they also shed light on potential mechanisms underlying this regulation by revealing the spatiotemporal dynamics of PI(3,4,5)P_3_ generation in living cells.

Regarding the biochemical experiments, an intriguing finding is that Es2 only weakly inhibits AKT phosphorylation in HeLa cells even though it inhibits both basal and EGF-stimulated PI(3,4,5)P_3_ levels in the same cells. Based on (i) the observed spatial confinement of PI(3,4,5)P_3_ generation by single exocytic events and (ii) a reported dependence of basal PI(3,4,5)P_3_ levels on the exocyst in a mammary epithelial cell line (NMuMG; Ahmed and Macara, 2017), we speculate that PI(3,4,5)P_3_ generation by exocytosis might be functionally coupled to AKT activation in epithelial cells but not in HeLa cells. This notion is supported by experiments showing that Es2 can strongly inhibit AKT stimulation in four different epithelial cell lines. The epithelial cell lines we tested not only originate from different tissues but also have different genetic backgrounds: HepG2 cells (liver) are nontumorigenic, whereas MCF7 (mammary) and DLD1 cells (colon) carry oncogenic mutations in *PIK3CA* and *KRAS*, respectively, and A431 cells (epidermis) overexpress EGFR. Although it remains to be determined whether non-epithelial cells, like HeLa cells, are generally less sensitive to Es2 with respect to AKT stimulation, we observed that when epithelial cells are not optimally cultured, under conditions in which the cells exhibit fibroblast-like morphology, they also show reduced sensitivity of AKT stimulation to Es2 (data not shown). Thus, the coupling between exocyst-mediated exocytosis and AKT stimulation is malleable and may depend on the physiological state of a cell.

An important aspect of our signaling experiments is that Es2 also inhibits MAPK stimulation, but at ∼3-fold higher concentrations. This is analogous to the differential sensitivity of AKT and MAPK stimulation to the EGFR TKI gefitinib. Thus, Es2 behaves like a low-affinity EGFR TKI. However, because Es2 does not inhibit EGFR auto-phosphorylation in MCF7 or A431 cells, it likely targets a signaling node that is downstream of EGFR activation but close to this initial signaling event. This idea is supported by the unexpected finding that AKT is reactivated during prolonged treatment with an EGFR TKI in A431 cells, and that this reactivation, which does not involve the relief of EGFR inhibition, can be negated by treating cells with Es2. Thus, it appears that exocyst-mediated exocytosis wholly subserves AKT stimulation by EGFR, in agreement with the ability of exocyst optogenetics to induce AKT stimulation. To our knowledge, this rapidly acquired resistance of AKT stimulation to EGFR TKIs has not been described before. Additional studies will be required to elucidate the upregulation of exocyst-mediated exocytosis in response to suppression of EGFR signaling, but it may involve the MAPK pathway as the reactivation of AKT stimulation does not occur when MAPK is completely inhibited (Fig. 5B).

Our model depicting the regulation of EGF-induced AKT signaling by exocyst-mediated exocytosis is presented in Figure 6. In the first step, EGFR signaling promotes exocytosis of recycling vesicles. It has been known that EGF can stimulate exocytosis of recycling vesicles for over two decades (Bretscher and Aguado-Velasco, 1998; Hopkins et al., 1994), but the mechanism underlying this process is still unclear. However, it may involve regulation of the exocyst, either by direct phosphorylation of Exo70 by MAPK to stabilize the exocyst complex (Ren and Guo, 2012), or through an interaction between the Rho family GTPase TC10 and Exo70 to promote binding of Exo70 to the plasma membrane (Inoue et al., 2003; Kawase et al., 2006). In a previous study (An et al., 2021), we showed that a vesicle needs to be tethered at the membrane by enough exocyst complexes for it to undergo full vesicle fusion; otherwise, the vesicle undergoes kiss-and-run fusion, which is a reversible mode of exocytosis that does not deliver vesicular cargo effectively. Thus, if EGF signaling promotes exocytosis by regulating vesicle tethering, it could do so by affecting the number of exocyst complexes that engage with the plasma membrane.

**Figure 6.**
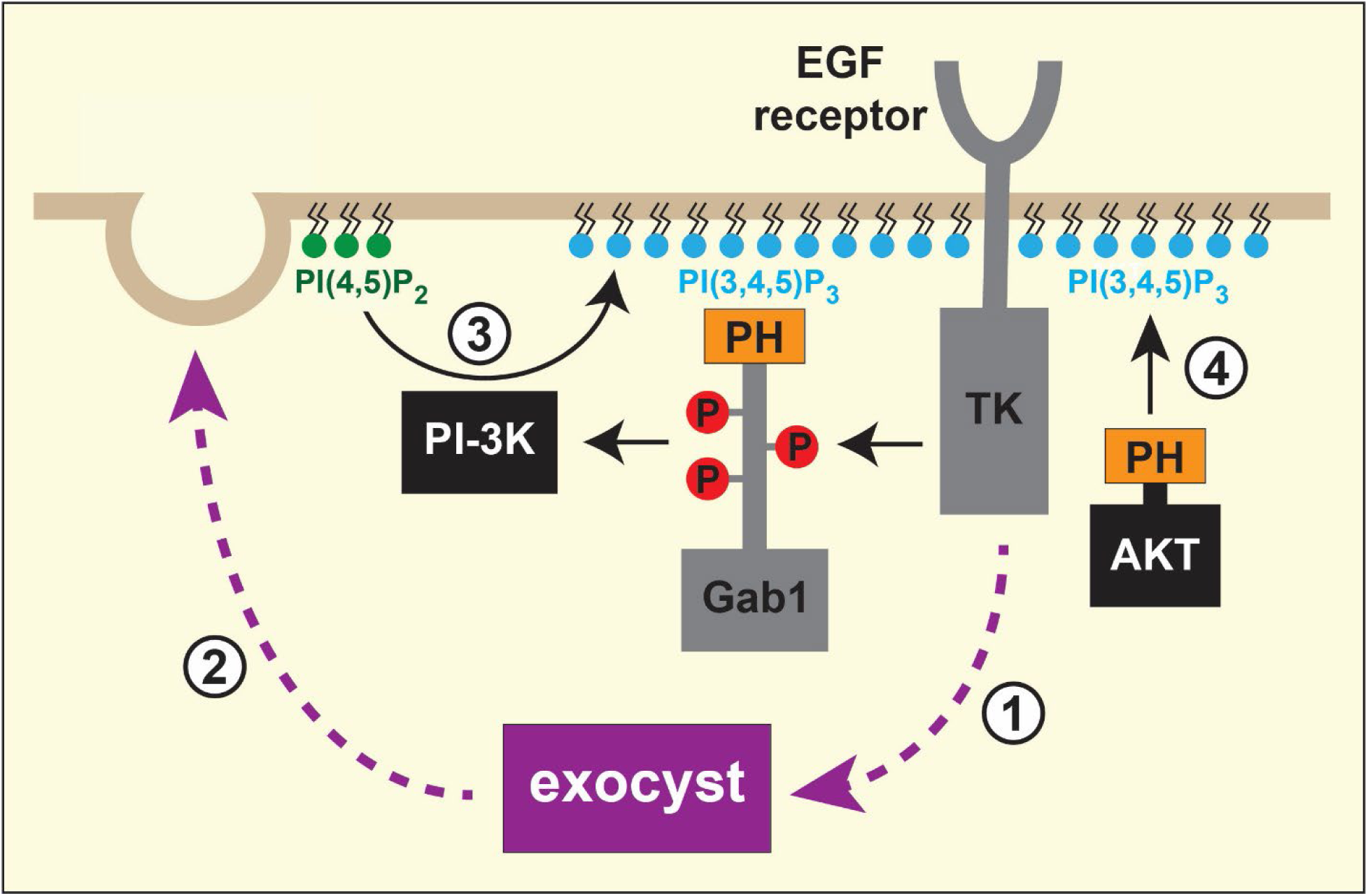
Regulation of EGF-stimulated activation of the PI-3K/AKT pathway by exocyst-mediated exocytosis in epithelial cells. (*Step 1*) EGF stimulation promotes exocyst-mediated tethering and exocytosis of recycling vesicles. (*Step 2*) Exocytosis delivers essential components or regulatory elements involved in PI(3,4,5)P synthesis. (*Step 3*) PI-3 kinase converts PI(4,5)P_2_ to PI(3,4,5)P_3_. (*Step 4*) AKT is recruited to the membrane via binding to PI(3,4,5)P_3_ for its activation. Note that Gab1 is also localized to the membrane by an indirect association with EGFR via Grb2 (not shown).

In the second step of the model, vesicles undergo exocytosis which in turn promotes PI(3,4,5)P_3_ synthesis at the membrane. While the mechanism underlying exocyst-induced PI(3,4,5)P_3_ production is not clear, we propose that essential components or regulatory elements are delivered to specific sites at the membrane to generate PI(3,4,5)P_3_. This hypothesis is based on the observation that the cloud of PI(3,4,5)P_3_ that forms after a single exocytic event is spatially confined to the fusion site for a relatively long time (∼1 min). Such local and persistent confinement of PI(3,4,5)P_3_ production suggests the presence of scaffolding proteins near the fusion site that couples the exocytic delivery of the unknown factors to PI(3,4,5)P_3_ synthesis. An attractive scaffolding candidate is the ubiquitously expressed IQ-motif-containing GTPase-activating protein 1 (IQGAP1), a multidomain protein that can associate with EGFR, the exocyst and components of several signaling pathways, including the PI-3K and MAPK pathways (Smith et al., 2015). Moreover, IQGAP1 has been shown to assemble all the phosphoinositide kinases – PI(4)KIIIα, PIPKIα (a.k.a. PI(4)P5K) and PI(4,5)P3K – that are necessary to sequentially convert phosphatidylinositol to PI(3,4,5)P_3_ (Choi et al., 2016). In this regard, it is interesting that recycling vesicles carry PI(4)P (Ketel et al., 2016) and that the exocyst complex interacts with PIPKIγi2 (Thapa et al., 2013). A scenario thus emerges whereby exocyst-mediated exocytosis could feed into the local synthesis of PI(4,5)P at the membrane to sustain PI(3,4,5)P_3_ levels.

In summary, we describe a novel role of the exocyst in EGFR signaling. We conclude that AKT activation by EGF in epithelial cells depends on the delivery of factors to the plasma membrane by exocyst-mediated exocytosis that promote the local generation of PI(3,4,5)P_3_. The scheme presented in Figure 6 depicts a model for how exocyst-mediated PI(3,4,5)P_3_ production will enhance both Gab1 and AKT recruitment to the cell membrane to facilitate a robust positive feedback mechanism for EGF stimulation of the PI-3K/AKT cascade in epithelial cells. Since exocytosis itself is promoted by EGF, AKT stimulation in epithelial cells thus involves the reciprocal regulation of EGFR signaling and exocytosis of recycling vesicles.

## Materials and Methods

### Plasmids and Reagents

To generate PH_AKT_-TagRFP-T plasmids, DNA sequences for wildtype and R25C mutant of PH_AKT_ from PH-Akt-GFP plasmid (#51465, Addgene) and PH-Akt(R25C)-GFP plasmid (#51466, Addgene) were amplified by PCR and cloned into pmTagRFP-T-N1 (Robert, Hermann, Davidson and Gelfand, 2014) using XhoI and BamHI sites. PH_AKT_-stagRFP was generated by site-directed mutagenesis of TagRFP-T (D159V) in the PH_AKT_-TagRFP-T plasmid. Exo70(K632A, K635A)-CRY2-mCherry, CIB1-CAAX and TfRc-pH were previously described (An et al., 2021). All cloning was done using standard molecular biology techniques. Endosidin2 (Es2) was purchased from Sigma or Cayman Chemical and stored as a 200 mM stock solution in DMSO at −30 °C. Wortmannin, gefitinib and afatinib were purchased from Tocris Bioscience and stored as 10 mM, 100 mM and 20 mM stock solutions, respectively, in DMSO at −30 °C.

### Cell culture and transfection

#### HeLa cells

For live-cell imaging experiments, HeLa cells (CCL-2, ATCC) were maintained in T-75 flasks (Falcon) at 37 °C and 5% CO_2_ in DMEM (Gibco) supplemented with 4.5 g/L glucose, 1 mM sodium pyruvate, 1× non-essential amino acids (Gibco), 10% (vol/vol) FBS (Sigma) and 100 U/ml penicillin-streptomycin mix (Gibco). Cells were split every 3−4 d, before they reached confluence. To transiently transfect HeLa cells for live-cell imaging experiments, a Nepa21 Type II electroporator (Nepa Gene) was used according to the manufacturer’s instructions. Briefly, 10^6^ cells were resuspended in ∼100 µl chilled Opti-MEM (Gibco) containing the following plasmids alone or in combination: 0.25 µg PH_AKT_-GFP, 0.25 µg PH_AKT_-TagRFP-T, 0.25 µg PH_AKT_-stagRFP, and 2 µg TfRc-pH. For exocyst optogenetics experiments, 10^6^ cells were resuspended in ∼100 µl chilled Opti-MEM (Gibco) containing 16 µg Exo70(K632A, K635A)-CRY2-mCherry, 4 µg CIB1-CAAX and 16 mM Exo70 siRNA (CCA UUG UGC GAC ACG ACU UTT; Sigma) and, if needed, 0.75 µg PH_AKT_-TagRFP-T and 2 µg TfRc-pH. Then, for all transfections, the cell mix was placed into a cuvette with a 2-mm electrode gap (EC-002) and subjected to sequential trains of two and five square-voltage pulses with the following settings, respectively: (i) 125 V, 3-ms pulse length, 50-ms pulse interval, 10% decay rate and (+) polarity and (ii) 25 V, 50-ms pulse length, 50-ms pulse interval, 40% decay rate and (±) polarity. Cells were then diluted into ∼12 ml phenol red-free DMEM (containing all supplements except antibiotics) and 2 ml aliquots were plated onto 35-mm glass-bottom dishes (MatTek). For exocyst optogenetics involving PI(3,4,5)P_3_ imaging, cells were cultured for 2−3 d before experiments to allow for molecular replacement of endogenous Exo70 with Exo70(K632A, K635A)-CRY2-mCherry. For whole-dish exocyst optogenetics experiments, transfected cells were plated on 35-mm treated polystyrene dishes (Falcon) and optically triggered with an LED box for 5 min at 37 °C and 5% CO_2_ as previously described (Xu et al., 2016).

For signaling experiments involving Es2, HeLa cells were maintained in T-75 flasks (Corning) at 37 °C and 5% CO_2_ in EMEM (ATCC) supplemented with 10% (vol/vol) γ-irradiated FBS (Corning). Cells were split at a 1:2 ratio every 2 d, using 1× TrypLE Express Enzyme (Gibco) to dissociate cells. For biochemical experiments, 2.5 × 10^5^ cells were plated onto 35-mm treated polystyrene dishes (Corning), cultured for 1 d, and then starved in serum-free media containing 0.2% BSA (Fraction V, low endotoxin grade; GeminiBio).

#### MCF7 cells

MCF7 cells (HTB-22, ATCC) were maintained in T-75 flasks (Corning) at 37 °C and 5% CO_2_ in EMEM (ATCC) supplemented with 0.01 mg/mL human recombinant insulin (Gibco) and 10% (vol/vol) γ-irradiated FBS (Corning). Cells were split at a 1:3 ratio every 4 d, using 1× TrypLE Express Enzyme (Gibco) to dissociate cells. For biochemical experiments, 1.875 × 10^5^ cells were plated onto 35-mm treated polystyrene dishes (Corning), cultured for the first 3 d in complete media, cultured for the next 4 d in media without insulin and then finally starved for 2 d in serum-free media containing 0.2% BSA (Fraction V, low endotoxin grade; GeminiBio).

#### A431 cells

A431 cells (CRL-1555, ATCC) were maintained in T-75 flasks (Corning) at 37 °C and 5% CO_2_ in DMEM (ATCC) supplemented with 10% (vol/vol) γ-irradiated FBS (Corning). Cells were split at a 1:2 ratio every 2 d, using 1× TrypLE Express Enzyme (Gibco) to dissociate cells. For biochemical experiments, 2.5 × 10^5^ cells were plated onto 35-mm treated polystyrene dishes (Corning), cultured for 3 d to 80% confluence and then starved for either 6 or 24 h in serum-free media containing 0.2% BSA (Fraction V, low endotoxin grade; GeminiBio).

#### DLD1 cells

DLD1 cells (CCL-221, ATCC) were maintained in T-75 flasks (Corning) at 37 °C and 5% CO_2_ in RPMI-1640 medium (ATCC) supplemented with 10% (vol/vol) γ-irradiated FBS (Corning). Cells were split at a 1:4 ratio every 2 d, using 1× TrypLE Express Enzyme (Gibco) to dissociate cells. For biochemical experiments, 5 × 10^5^ cells were plated onto 35-mm treated polystyrene dishes (Corning), cultured for 4 d to 100% confluence and then starved for either 6 or 24 h in serum-free media containing 0.2% BSA (Fraction V, low endotoxin grade; GeminiBio).

#### HepG2 cells

HepG2 cells (HB-8065, ATCC) were maintained in T-75 flasks (Corning) at 37 °C and 5% CO_2_ in EMEM (ATCC) supplemented with 10% (vol/vol) γ-irradiated FBS (Corning). Cells were split at a 1:4 ratio every 2 d, using 1× TrypLE Express Enzyme (Gibco) to dissociate cells. For biochemical experiments, 1.25 × 10^5^ cells were plated onto 35-mm treated polystyrene dishes (Corning), cultured for 3 d to 80% confluence and then starved for either 6 or 24 h in serum-free media containing 0.2% BSA (Fraction V, low endotoxin grade; GeminiBio).

### Total internal reflection fluorescence microscopy

Live-cell imaging was done as previously described (An et al., 2021) using a custom TIRF microscope equipped with 488- and 568-nm solid state lasers (Melles Griot), an EMCCD camera and a 60× 1.49 NA TIRF objective (Olympus). All experiments were done at 37 °C (using a custom incubator chamber) in phenol red-free DMEM with 25 mM HEPES, pH 7.4, with or without 10% FBS. Image frames were acquired in time-lapse recordings at 2 or 0.2 Hz with 150-ms exposures. Pixel size was 160 nm. For PI(3,4,5)P_3_ imaging with PH_AKT_-stagRFP, cells were imaged at 37 °C and 5% CO2 in a cage incubator (OkoLab) housing a Nikon Eclipse Ti2 microscope (Nikon) equipped with a motorized Ti-LA-HTIRF module with a 15-mW LU-N4 488- and 561-nm lasers, using a CFI Plan Apochromat Lambda 100×/1.45 Oil TIRF objective and a Prime95B cMOS camera (110-nm pixel size; Teledyne Photometrics). Images were acquired using a 100-ms exposure time at 1 Hz. Pixel size was 160 nm.

### Immunoprecipitation and immunoblotting

For immunoprecipitation, cells were lysed using lysis buffer (50 mM HEPES, pH 7.5, 150 mM NaCl, 1mM EDTA, 1mM EGTA, 1% Triton X-100, 10% glycerol, 25mM NaF, 1mM MgCl2, 1mM sodium orthovanadate (Na_3_VO_4_), and complete protease inhibitor cocktail (Roche)) and lysates were clarified by centrifugation at 13,000 g. Protein A/G Agarose beads (Thermo Fisher Scientific) and Anti-EGFR antibody EGFR (100:1 dilution, D38B1; Cell Signaling Technology, #4267) were added directly to the cell lysate. After overnight incubation of antibodies, beads and lysate at 4 °C with shaking, immune complexes were collected and washed three times with 1 mL washing buffer (20mM HEPES, 150mM NaCl, 1% Triton X-100, 5% glycerol) at 4 °C. 2× loading buffer was added and samples were boiled at 95 °C for 5 min, spun to pellet the beads and subjected to SDS–PAGE. Western blot analysis was performed using antibodies specific to EGFR (D38B1; Cell Signaling Technology, #4267), pEGFR (53A5, Tyr1173; Cell Signaling Technology, #4407), AKT (Cell Signaling Technology, #9272), pAKT (D9E, Ser473; Cell Signaling Technology, #4060), pMAPK (Thr202/Tyr204; Cell Signaling Technology, #9101), and γ-tubulin (GTU-88; Sigma, #T6557). Immunoreactive bands were detected with horseradish peroxidase-conjugated secondary antibodies through chemiluminescence.

## Acknowledgments

Supported by grants from the National Institutes of Health (R01GM098498 and R01GM118486) and the Wellcome Trust Foundation (095927/A/11/Z).

