## Supplementary Fig. 1 for "Regulation of EGF-stimulated activation of the PI-3K/AKT pathway by exocyst-mediated exocytosis"

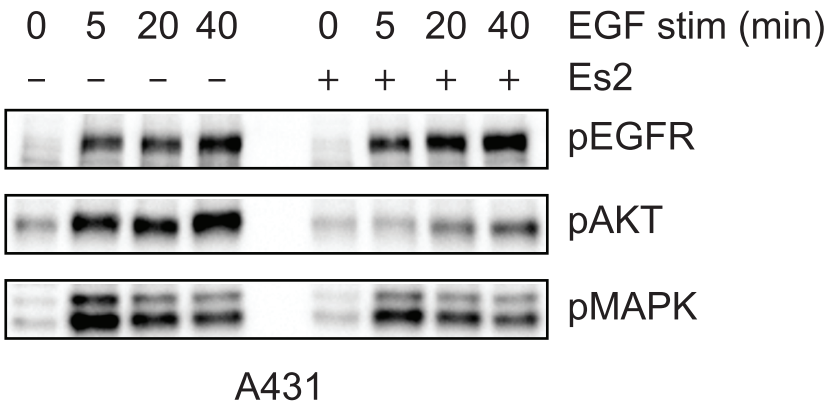


**Supplementary Figure 1. Es2 inhibits AKT stimulation but not EGFR activation in A431 cells**

A431 cells were cultured for 3 d, starved for either 6, treated with Es2 (240 μM) for 30 min at 37 °C, and then stimulated with EGF (10 ng/mL) for 5 min. Cell lysates were subjected to SDS/PAGE and analyzed for activation of EGFR, AKT and MAPK by immunoblotting with antibodies for pEGFR (Tyr 1068), pAKT (Ser473) and pMAPK (Thr202, Tyr204), respectively.
